# Preparing for the next COVID: Deep Reinforcement Learning trained Artificial Intelligence discovery of multi-modal immunomodulatory control of systemic inflammation in the absence of effective anti-microbials

**DOI:** 10.1101/2022.02.17.480940

**Authors:** Dale Larie, Gary An, Chase Cockrell

## Abstract

**Background:** Despite a great deal of interest in the application of artificial intelligence (AI) to sepsis/critical illness, most current approaches are limited in their potential impact: prediction models do not (and cannot) address the lack of effective therapeutics and current approaches to enhancing the treatment of sepsis focus on optimizing the application of existing interventions, and thus cannot address the development of new treatment options/modalities. The inability to test new therapeutic applications was highlighted by the generally unsatisfactory results from drug repurposing efforts in COVID-19.

**Hypothesis:** Addressing this challenge requires the application of simulation-based, model-free deep reinforcement learning (DRL) in a fashion akin to training the game-playing AIs. We have previously demonstrated the potential of this method in the context of bacterial sepsis in which the microbial infection is responsive to antibiotic therapy. The current work addresses the control problem of multi-modal, adaptive immunomodulation in the circumstance where there is no effective anti-pathogen therapy (e.g., in a novel viral pandemic or in the face of resistant microbes).

**Methods:** This is a proof-of-concept study that determines the controllability of sepsis without the ability to pharmacologically suppress the pathogen. We use as a surrogate system a previously validated agent-based model, the Innate Immune Response Agent-based Model (IIRABM), for control discovery using DRL. The DRL algorithm ‘trains’ an AI on simulations of infection where both the control and observation spaces are limited to operating upon the defined immune mediators included in the IIRABM (a total of 11). Policies were learned using the Deep Deterministic Policy Gradient approach, with the objective function being a return to baseline system health.

**Results:** DRL trained an AI policy that improved system mortality from 85% to 10.4%. Control actions affected every one of the 11 targetable cytokines and could be divided into those with static/unchanging controls and those with variable/adaptive controls. Adaptive controls primarily targeted 3 different aspects of the immune response: 2^nd^ order pro-inflammation governing TH1/TH2 balance, primary anti-inflammation, and inflammatory cell proliferation.

**Discussion:** The current treatment of sepsis is hampered by limitations in therapeutic options able to affect the biology of sepsis. This is heightened in circumstances where no effective antimicrobials exist, as was the case for COVID-19. Current AI methods are intrinsically unable to address this problem; doing so requires training AIs in contexts that fully represent the counterfactual space of potential treatments. The synthetic data needed for this task is only possible through the use of high-resolution, mechanism-based simulations. Finally, being able to treat sepsis will require a reorientation as to the sensing and actuating requirements needed to develop these simulations and bring them to the bedside.

## Introduction

While the impact of the COVID-19 pandemic is ongoing and the full story of the pandemic is yet to be known one thing that is highly likely is that the future historical consideration of the pandemic will invariably focus on the early phases of the pandemic when medical resources, particularly those in critical care units, were overwhelmed. A notable issue during that time (and, to a great degree, continues) was the inability to affect the underlying processes that drove the course of disease; once the disease manifested the only option was supportive care until the disease ran its course. There were numerous efforts to develop interventions that could potentially affect the biology of COVID, including repurposing of existing drugs. Specifically, there was a relatively early recognition that severe disease was associated with “cytokine storm” (1–6), namely that the body’s inflammatory/immune response was producing unintended and detrimental collateral damage in response to the viral infection. As a result, there was a great deal of interest in repurposing immunomodulatory agents to attempt to mitigate disease severity (7–9), but to date, with the exception of the use of steroids for severe disease (10), none of these approaches have been proven to be effective.

This should not come as a surprise. The phenomenon of collateral tissue damage arising from dysregulated inflammation described as “cytokine storm” is exactly the process that drives disease severity and multiple organ failure in bacterial sepsis, for which no immunomodulatory interventions have been shown to be effective (11). In fact, the current set of immunotherapies for chronic inflammatory diseases, exactly those proposed for use in COVID, were repurposed from agents that initially failed in sepsis trials. We have previously reported on the challenges present in attempting to control sepsis using anti-cytokine/anti-mediator therapies, primarily stemming from the failures to recognize the dynamic complexity of the mechanistic processes ostensibly being targeted (12) and that in order to be effective the treatment of sepsis should be considered a complex control problem (13). In previous work we have shown that clinical sepsis is controllable using different types of ML methods for control discovery on the IIRABM (14, 15). Specifically, the latter project utilized the same method, DRL, as has been used by successful ML/Artificial Intelligence (AI) systems, the game-playing AIs and their descendants from Deep Mind (16–20). We term this approach simulation-based, model-free DRL. In our prior work we treated the attempt to control sepsis as a “game” to be played using the IIRABM, where potential cytokine interventions represented the “moves” implemented by the AI agent (15). However, those investigations attempted to mimic the current standard of care for the treatment of bacterial sepsis and therefore included the administration of antibiotics that could directly reduce the microbial load. There are no effective antiviral agents for COVID-19 (as well as the vast majority of acute viral illnesses), and, furthermore, there is a suggestion that given the time courses of the pathogenesis of acute viral infections (1–6) disease manifestation occurs subsequent to the peak(s) of viremia. We propose that given:

1. Dysregulated and detrimental systemic inflammation is a primary source of disease severity in acute viral illness;
2. There is a critical need to have virus-agnostic disease mitigation therapies in the early phases of a pandemic; and
3. There is proven inefficacy of standard approaches to applying immunomodulation in the face of cytokine storm/sepsis,

the application of simulation-based control discovery using DRL can provide useful insights and potentially critical capabilities in designing effective multi-modal and adaptive immunomodulatory therapies for infections for which no effective anti-microbial agents exist. Herein we present the application of DRL to train an artificial neural network (ANN) to discover a treatment policy to improve the outcomes to simulated infection in the absence of anti-microbial treatment, with the observation space limited to clinically measurable observables.

## Methods

### Description of IIRABM

The simulation model in this work is a previously validated agent-based model of sepsis, the Innate Immune Response agent-based model (IIRABM) (12, 21). We have previously used the IIRABM as a surrogate/proxy system for the investigation of potential control strategies (22) for sepsis, both using genetic algorithms (14) and DRL (15). The IIRABM is a two-dimensional abstract representation of the human endothelial-blood interface with the modeling assumption that the endothelial-blood interface is the initiation site for acute inflammation. The closed nature of the circulatory surface can be represented as a torus, and the two-dimensional surface of the IIRABM therefore represents the sum-total of the capillary beds in the body. The spatial scale of the real-world system is not directly mapped using this scheme. The IIRABM simulates the cellular inflammatory signaling network response to injury/infection and reproduces all the overall clinical trajectories of sepsis (21). The IIRABM incorporates multiple cell types and their interactions: endothelial cells, macrophages, neutrophils, TH0, TH1, and TH2 cells as well as their associated precursor immune cells. This is not intended to be a comprehensive list of all the cellular subtypes present in the immune system, but rather represents the minimally sufficient set of cell populations able to represent every necessary function in the innate response to infection. System mortality of the IIRABM is defined when the aggregate endothelial cell damage) exceeds 80%; this threshold represents the ability of current medical technologies to keep patients alive (i.e., through organ support machines) in conditions that previously would have been lethal. Infectious insults to the IIRABM are initiated using 5 parameters representing the size and nature of the injury/infection as well as a metric of the host’s resilience: initial injury size, microbial invasiveness, microbial toxigenesis, environmental toxicity, and host resilience. Previous work (12) identified the boundary conditions for these parameters in terms of generating clinically realistic behavior, and therefore we consider this parameter space as representing the clinically plausible space of human response to infection.

### Deep Reinforcement Learning

Deep Deterministic Policy Gradient (DDPG) (23) was used to discover a controls algorithm that is able to heal *in silico* patients by either augmenting or diminishing the concentration of cytokine signaling molecules in the simulation. DDPG is a powerful reinforcement learning (RL) algorithm able to use off-policy data and the Bellman equation (Equation 1) to learn a value function, or Q-function, to determine the most valuable action to take given any particular state of the simulation.

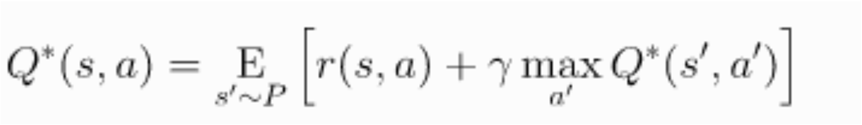

Equation 1: The Bellman Equation. Value *Q* is a function of the current state and action *(s, a)*, and is equal to the reward *r* from the current state and chosen action *(s, a)* summed with the discounted value of the next state (discount factor = γ) and action *(s’, a’)* where the next state is sampled from a probability distribution *(s’ ∼ P)*.

The Q-function is discovered through trial and error and allowing an RL agent to optimize the Q-function based on observed rewards from chosen actions. DDPG can be thought of as an extension of the Q-learning algorithm (24), where it is able to choose from a continuous action space. In Q-learning, the next action is chosen from a set of discrete actions that can be taken based on the output of the Q-function. The best action to take from the current state is identified by finding which action will return the highest value from the Q-function. Q-learning is an off-policy algorithm, which means that in the training phase, the RL agent is sometimes able to choose actions that are not the ones chosen by the Q-function. This allows the agent to explore and potentially discover actions that can lead to a greater reward than continuing from an already discovered policy. Q-learning has proven to be very powerful at solving control problems in discrete space and has proven on benchmark RL problems that the discovered controls algorithm can be very robust (23).

DDPG extends Q-learning to a continuous action space. It is too computationally expensive to exhaustively search the action space for the optimal action during the learning phase since the action space is continuous. Because of this, DDPG uses an “actor” neural network to choose an action based on the current state. The chosen action is used for the simulation, and a new state and reward is returned by the RL environment. The reward then updates the Q-function to more closely approximate the true value function for the environment, and the updated Q-function is used to perform gradient descent on the actor network to improve its decision making in the future. Because updates to the actor network are made based on an approximation, DDPG is sometimes susceptible to starting conditions, and is sometimes unstable as it learns. Because of this, learning rates for the actor network and the Q-function approximation are usually slow. Additionally, to help stability, DDPG uses what is called an “Experience Replay Buffer” to sample a batch of states and actions the agent has taken in the past, instead of relying only on the current state and action for network updates.

### Training Environment

The goal of this work is to determine if an effective immunomodulatory strategy would be successful in balancing the need for an effective immune response to contain an infection in the absence of anti-microbials while preventing system death due to cytokine storm. For purposes of evaluating this possibility we chose a high-mortality condition with a relatively virulent microbe. Using our previously identified method of finding relevant parameter sets within bioplausible parameter space (12) we chose the following External parameters and initial infection level:

- Host Resilience [`oxyheal`] = 0.08: This represents the rate at which the baseline endothelial cells recover their oxy level, back to a baseline of 100.
- Invasiveness [`infectSpread`] = 4: This represents the number of adjacent grid spaces the infection spreads to after it has reaches the carrying capacity on an individual grid.
- Environmental Toxicity [`numRecurInj`] = 2: This represents the number of grid spaces are randomly reinfected every 24 hours, reflecting environmental contamination.
- Toxigenesis [`numInfectRep`] = 2: This represents the amount of damage produced by a microbe on the grid space it occupies.
- Initial Infection Amount [`inj_number`] = 27: This represents the radius in number of grid spaces of a circular inoculation of the infection

With these parameters, the mortality of the IIRABM = 85% (15% Completely Healed).

#### Initial and Termination Conditions

An episode begins 12 hours after the application of the initial infection; this is to reflect the minimal necessary incubation time between exposure and initiation of any treatment. The episode ends when either the simulated patient completely heals, dies, or 10,000 time steps (= 42 days simulated time) if neither of those conditions is met.

#### Observation Space

The IIRABM states exists over a discrete, 2-dimensional 101 × 101 grid. The IIRABM includes 9 cytokines, 2 soluble cytokine receptors (essentially inhibitors of their respective cytokines), population levels of 5 different cell types, the total amount of infection in the system and the total amount of damage present in the system (as reflected by the variable “`Oxy-deficit`”). Since the IIRABM utilizes an abstract spatial representation, the individual discrete grids are not directly translatable to any potential measurement. Rather, the aggregated system levels are considered equivalent to values potentially sampled in the blood, and therefore represent the accessible information for any potential control-sensor. As this is a proof-of-concept investigation, we assume that any circulating cytokine/soluble receptor can be measured at every time step (= 6 minutes): this gives the system state as reflected in 11-dimensions (e.g. 9 cytokines + 2 soluble receptors represented in the IIRABM, hereafter termed “cytokines”). Also, since the total amount of damage in the system (“`Oxy-deficit`”) is not actually a quantifiable or observable metric in the clinical patient, this value is not included in the observations used to train the DRL (this is in contrast to our prior use of DRL trained on the IIRABM (15)); as such the current DRL agent is being trained on partially-observable states of the IIRABM.

#### Action space

The actions taken by the DRL agent can either be supplementation or inhibition of any or all of the 11 cytokines represented in the IIRABM every time step (= 6 minutes). Supplementation takes the form of the addition of a continuous value from 1 to 10 to the value of a particular cytokine. Inhibition can take the form of the multiplication of the existing cytokine value by 0.001 to 1; this approach is done to avoid negative (or exploding, in the case of pathway augmentation) values and is consistent with the dynamics of mediator inhibition. These are reflected in the code thusly:

`if action_mag > 0, action = (action_mag*9) +1` ⇒ add cytokine between 1 and 10

`If action_mag <0, action = action_mag + 1.001` ⇒ multiply cytokine between .001 and 1

The ability to manipulate any combination of cytokines present is meant to simulate the potential use of combinations of interventions, and the DRL is intended to assist in addressing the exponential combinatorial issues associated with multi-drug therapy.

#### Reward function

The current work includes two types of reward functions. The first are considered terminal rewards: these are evaluated at the end of an episode and are analogous to either winning or losing the game. The current work has a positive terminal reward if the system heals: *r* = 0.999^*step*^ * 1000, whereas the negative terminal rewards if the system dies is: *r* = 0.999^*step*^ − 1000. The incorporation of the step at which the terminating condition is met is intended to reward quicker healing, penalize faster death, and not penalize prolongation of life (albeit in a diseased state). The current work also includes intermediate or step-wise rewards; these are reinforcing conditions to aid in learning during the course of the episode run. The intermediate reward function is:

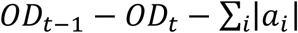

Where *OD*_*t*_ indicates the oxygen-deficit at time *t*, and *a*_*i*_ is the value for the action taken on mediator *i*. This calculation rewards systems that reduce their damage per time step and are able to do so with a minimal amount of intervention. The latter goal is consistent with the concept of minimizing necessary interventions and avoiding potential side effects that may not be reflected in the resolution of the simulation.

Code for the DRL environment can be found at https://github.com/An-Cockrell/DRL_Control

## Results

DRL training proceeded for 400 episodes and converged to a policy that had a Post Control Complete Healing/Recovered rate = 89.6% and a “Timeout” group of 10.4%, where this last group were system runs that never met the threshold for “death” but were unable to recover for the max simulation period of ∼ 42 days. We consider this “Timeout” group as the equivalent of Death, as once the control is released these systems die immediately (n = 500, with 448 Complete Healing/Recovered and 52 Timeouts/Deaths). These results were a significant improvement over the uncontrolled base condition, which had a Mortality of 85% (= 15% Complete Healing/Recovered). The discovered policy was also able to eradicate both the initial infection and subsequent reinfections without the aid of antibiotics.

A plot of the `Oxy-Deficit` (= total system damage) trajectories for both Recovered and Timeout/Death system runs can be seen in Figure 1.

**Figure 1:**
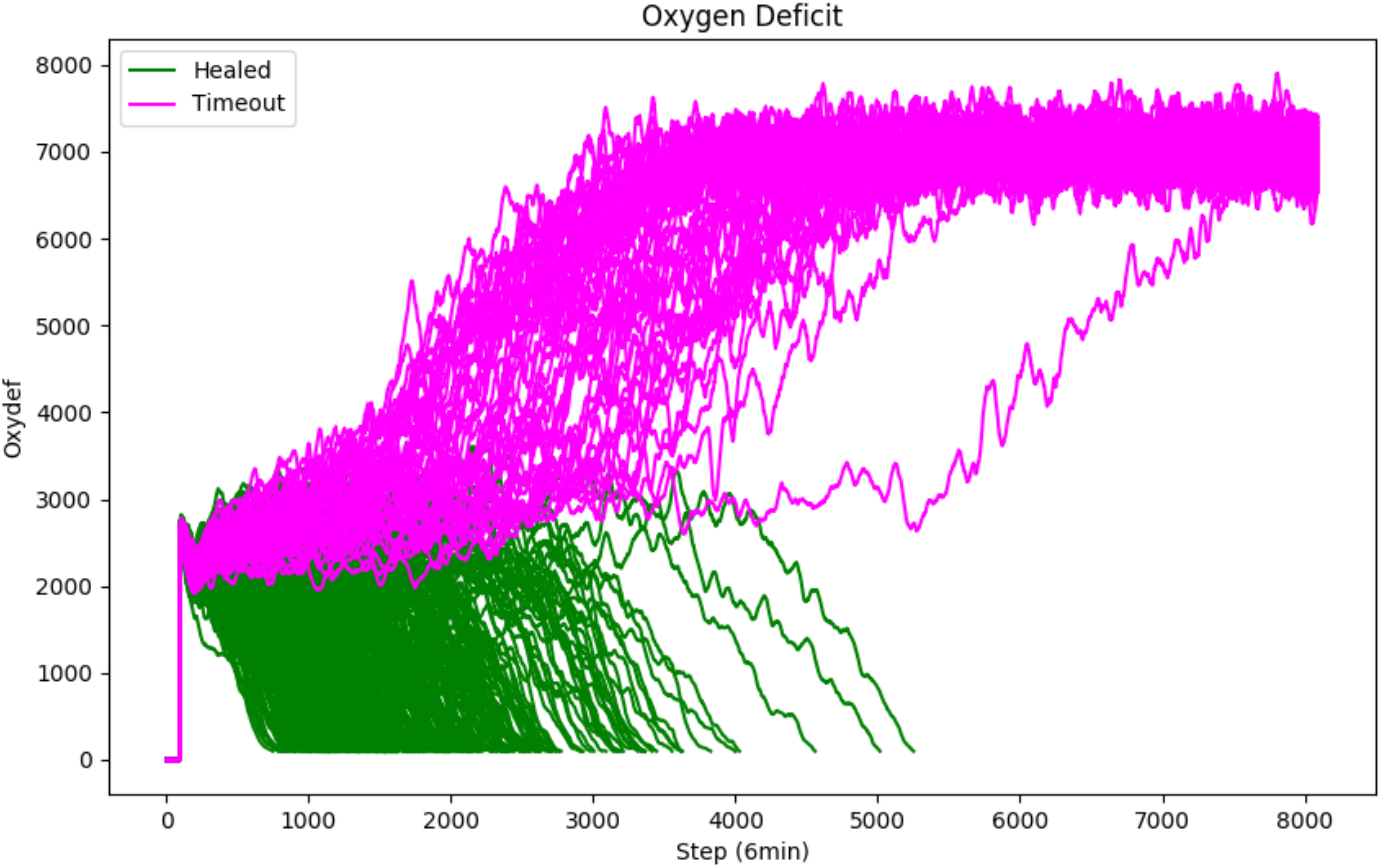
Oxy-Deficit (= Total System Damage) Trajectories with DRL control policy. N = 500, Completely Healed/Recovered (Green) = 448, Timeouts/Death (Pink) = 52

Of note, there appears to be a threshold of ∼ `4000 Oxy-deficit` at which the system cannot be steered back towards recovery.

In terms of evaluating the components of the discovered policy it is notable that every cytokine was manipulated in some fashion, therefore the policy involves coordinating 11 distinct interventions every time step. The overall control policy includes relatively static (unchanging) actions, as well as a series of cytokines that are subject to interventions that vary both in time and in magnitude. The cytokines that had static policies can be subdivided into those that essentially maximal augmentation for the duration of the runs and those that had maximal inhibition for the duration of the runs. Those with maximal augmentation include: Platelet Activating Factor (PAF) and Soluble Tumor Necrosis Factor Receptor (sTNFr), whereas those with maximal inhibition includes Tumor Necrosis Factor (TNF).

In terms of targeted cytokines that have varying manipulations, this group can be further divided into those that had variable intervention during the period of the primary/initial infection, and then settled into a static policy after the infection was eradicated (where the resulting static policy was maximal augmentation). Cytokines that fell into this group include: Interleukin-1 (IL1) (Figure 2) and Interleukin-4 (IL4) (Figure 3). *Note that in the following figures the Green-Recovered trajectories (= 448) vastly outnumber the Pink-Timeout/Death trajectories (= 52).

**Figure 2:**
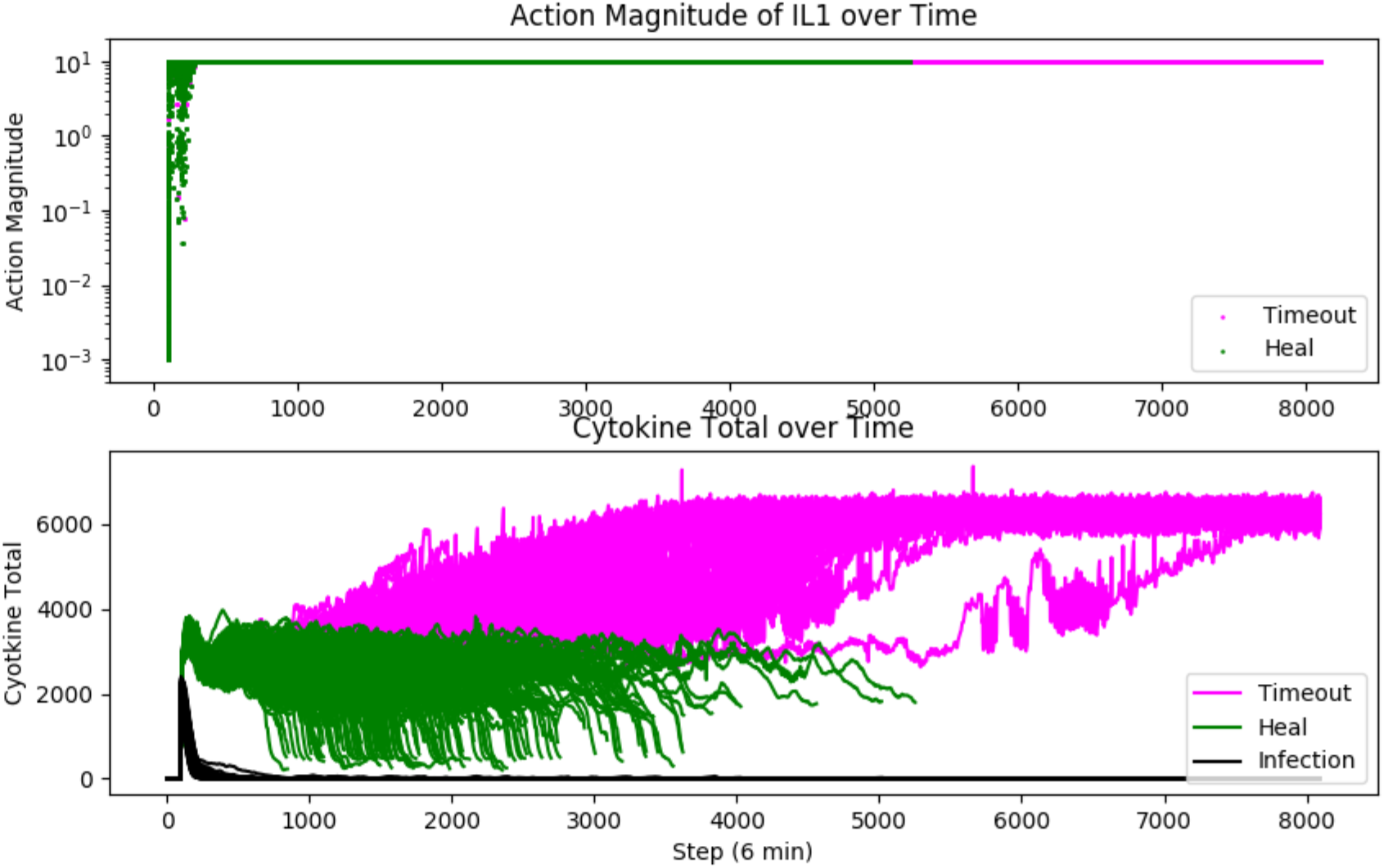
Upper panel: Action Magnitude of Control Policy targeting IL1. Lower panel: Controlled level of total system cytokines. Green = Recovered Simulations, Pink = Timeout/Death Simulations, Black Line Lower Panel = Level of Infection. Variable levels of control can be seen in the Upper Panel corresponding to the period during which the infection is being controlled.

**Figure 3:**
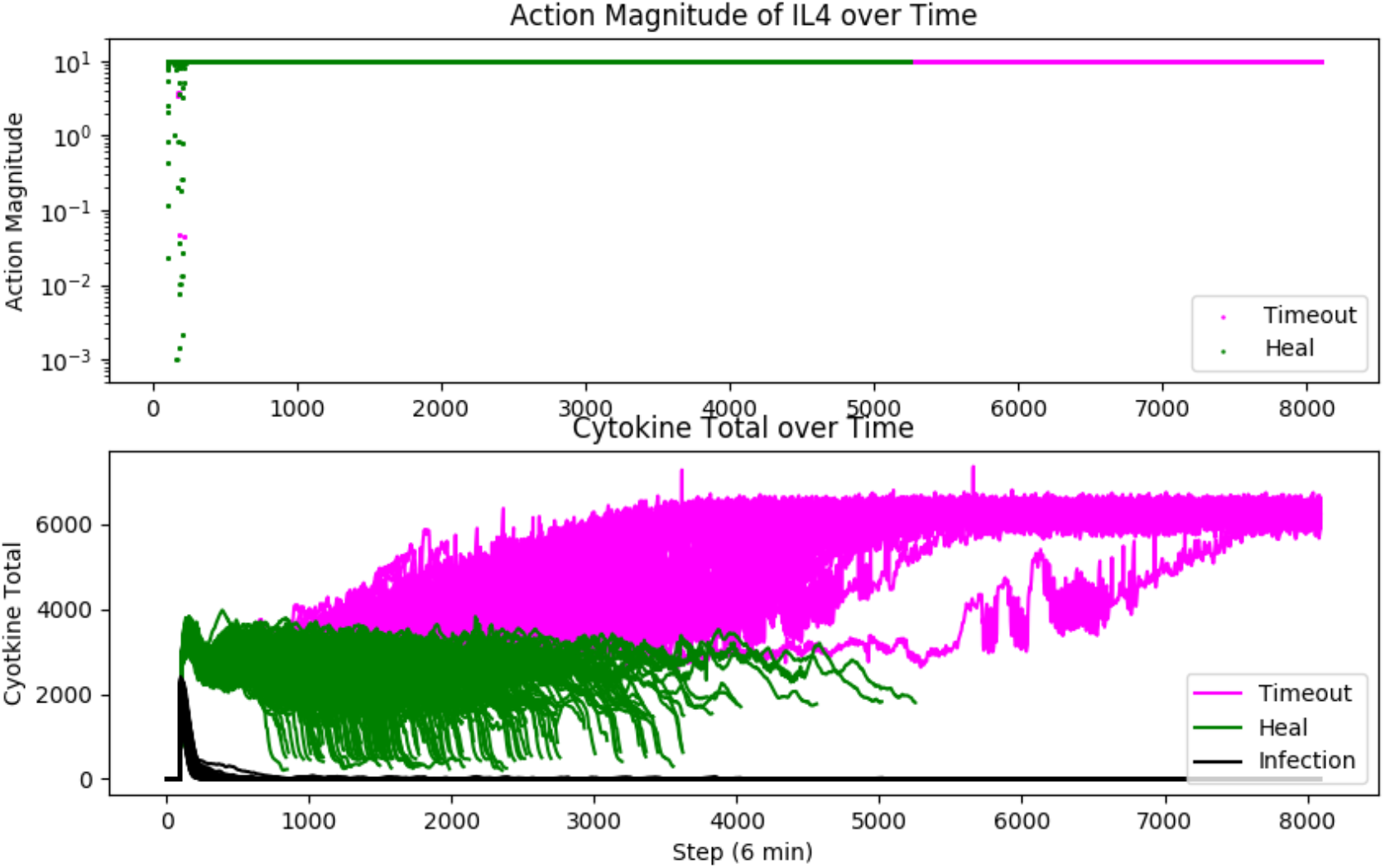
Upper panel: Action Magnitude of Control Policy targeting IL4. Lower panel: Controlled level of total system cytokines. Green = Recovered Simulations, Pink = Timeout/Death Simulations, Black Line Lower Panel = Level of Infection. Variable levels of control can be seen in the Upper Panel corresponding to the period during which the infection is being controlled.

Note that the Green/Recovered trajectories terminate before the total allowable simulation period, denoting that they have recovered. Also note that the Timeout/Death simulations manifest persistently elevated cytokine levels even despite maximal control.

The remaining controllable cytokines show a greater degree of variability, both within the successful, Recovered population and in comparison between the Recovered and Timeout/Death populations. These targets can be grouped into representing different phases of the initial phase of injury:

- 2^nd^ order pro-inflammation, affecting T-cell level determinants of TH1/TH2 balance, represented by Interferon-γ or IFNg (Figure 4)
- Anti-inflammation/pro-growth, represented by Interleukin-10 or IL10 (Figure 5)
- Inflammatory cell proliferation, represented by Granulocyte Colony Stimulating Factor or GCSF (Figure 6).

**Figure 4:**
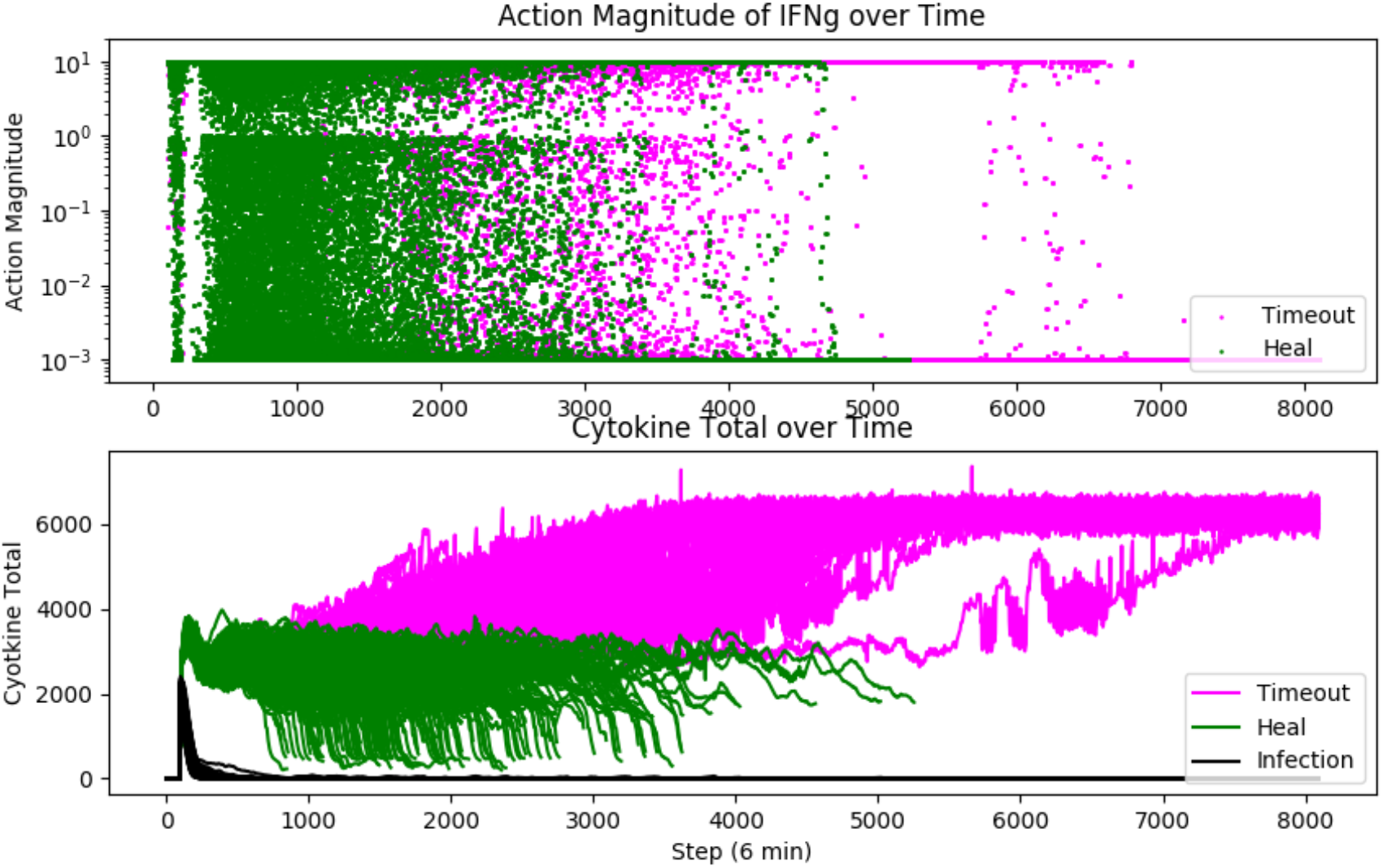
Upper panel: Action Magnitude of Control Policy targeting IFNg. Lower panel: Controlled level of total system cytokines. Green = Recovered Simulations, Pink = Timeout/Death Simulations, Black Line Lower Panel = Level of Infection. Note that the actions implemented are highly variable (as reflected by the scatter of points along the y-axis in the upper panel) throughout the entire simulation run for Recovered simulations. A similar variability is seen in the Timeout simulations.

**Figure 5:**
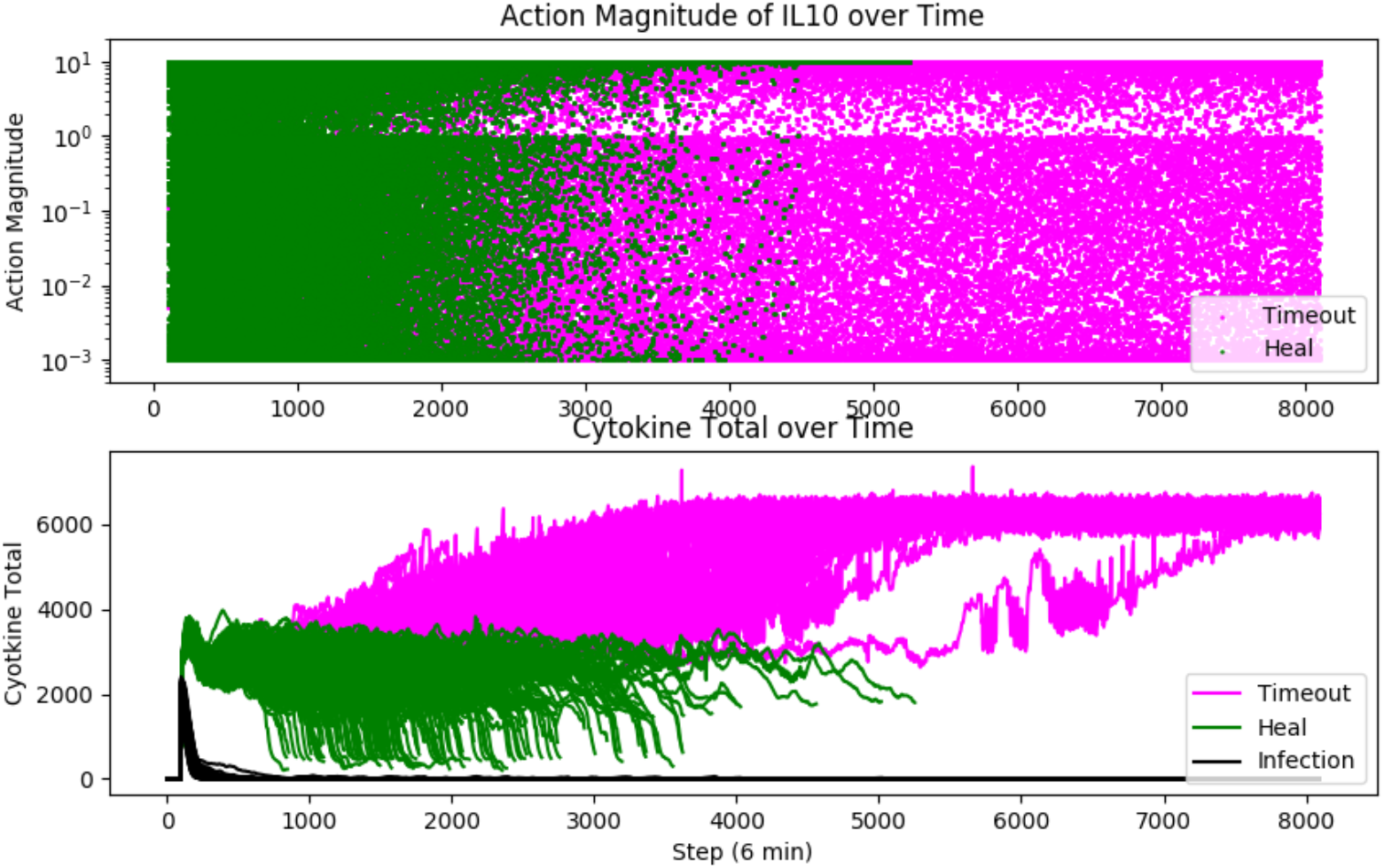
Upper panel: Action Magnitude of Control Policy targeting IL10. Lower panel: Controlled level of total system cytokines. Green = Recovered Simulations, Pink = Timeout/Death Simulations, Black Line Lower Panel = Level of Infection. Note that the actions implemented are highly variable (as reflected by the scatter of points along the y-axis in the upper panel) compared to actions taken in controlling IFNg seen in Figure 4.

**Figure 6:**
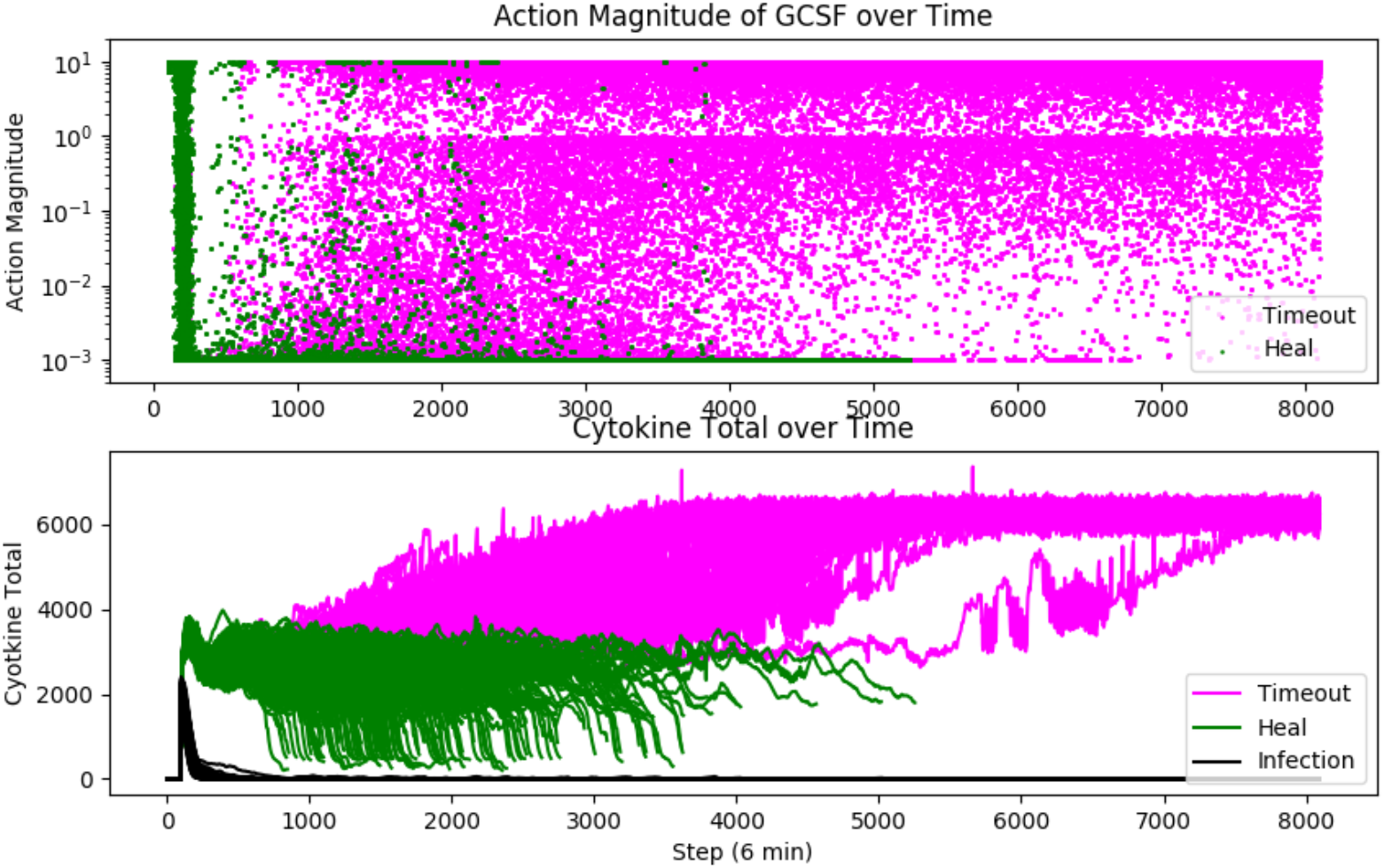
Upper panel: Action Magnitude of Control Policy targeting GCSF. Lower panel: Controlled level of total system cytokines. Green = Recovered Simulations, Pink = Timeout/Death Simulations, Black Line Lower Panel = Level of Infection. Note that the actions controlling GCSF in the Recovered simulations trend towards inhibition, particularly after the infection is eradicated, whereas actions implemented in the Timeout simulations remain highly variable.

To further visualize the effect of the various controls, the following Figures 7–9 demonstrate the “target” level of the manipulated cytokine as it is controlled, as well as the relationship between the degree of control and the targeted cytokine.

**Figure 7:**
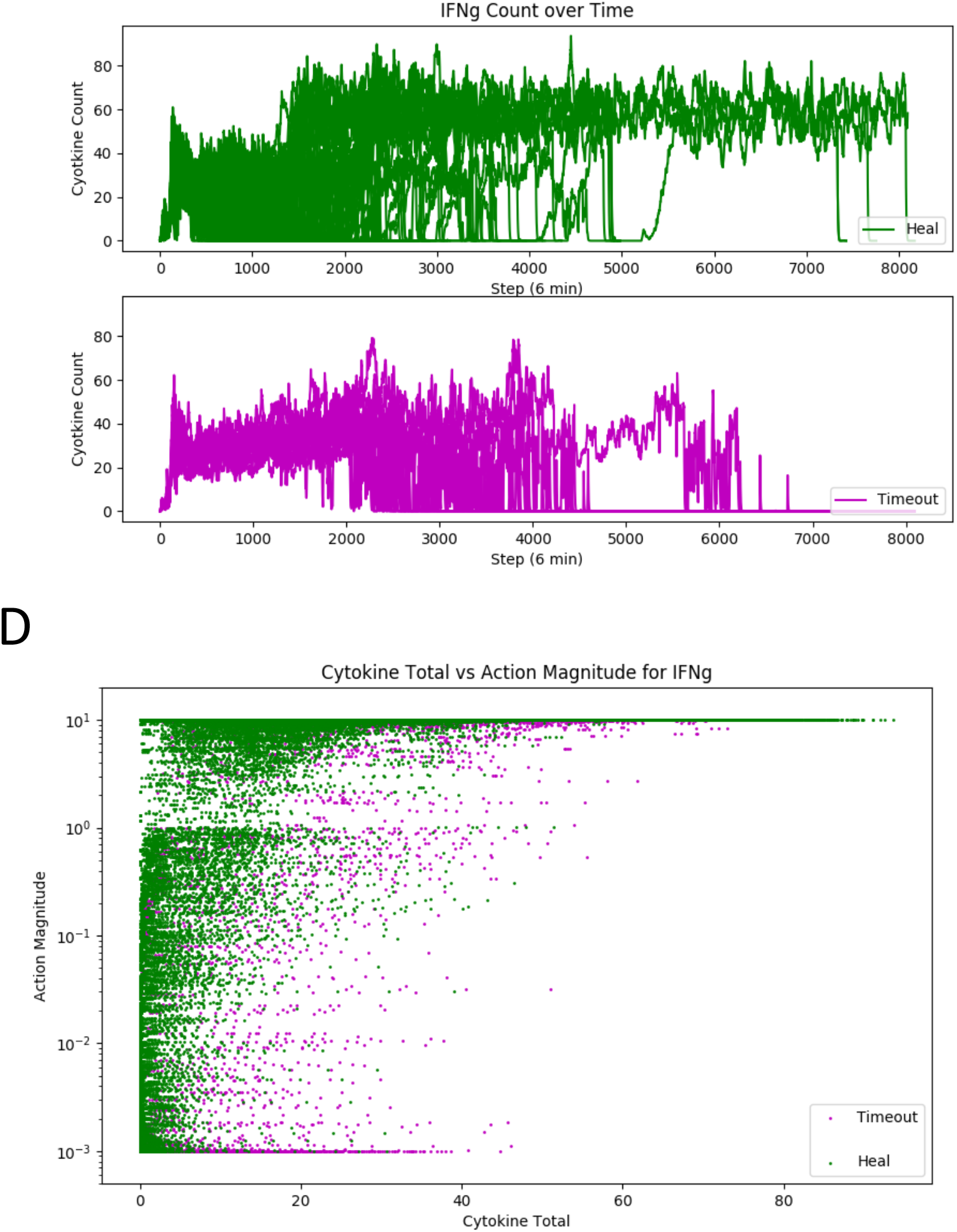
Upper panel: “Controlled” level of IFNg in Recovered simulations. Middle Panel: “Controlled” level of IFNg in Timeout/Death simulations. Lower Panel: Plot of Action Magnitude (= Control) on Y axis versus Target IFNg level on X-axis.

**Figure 8:**
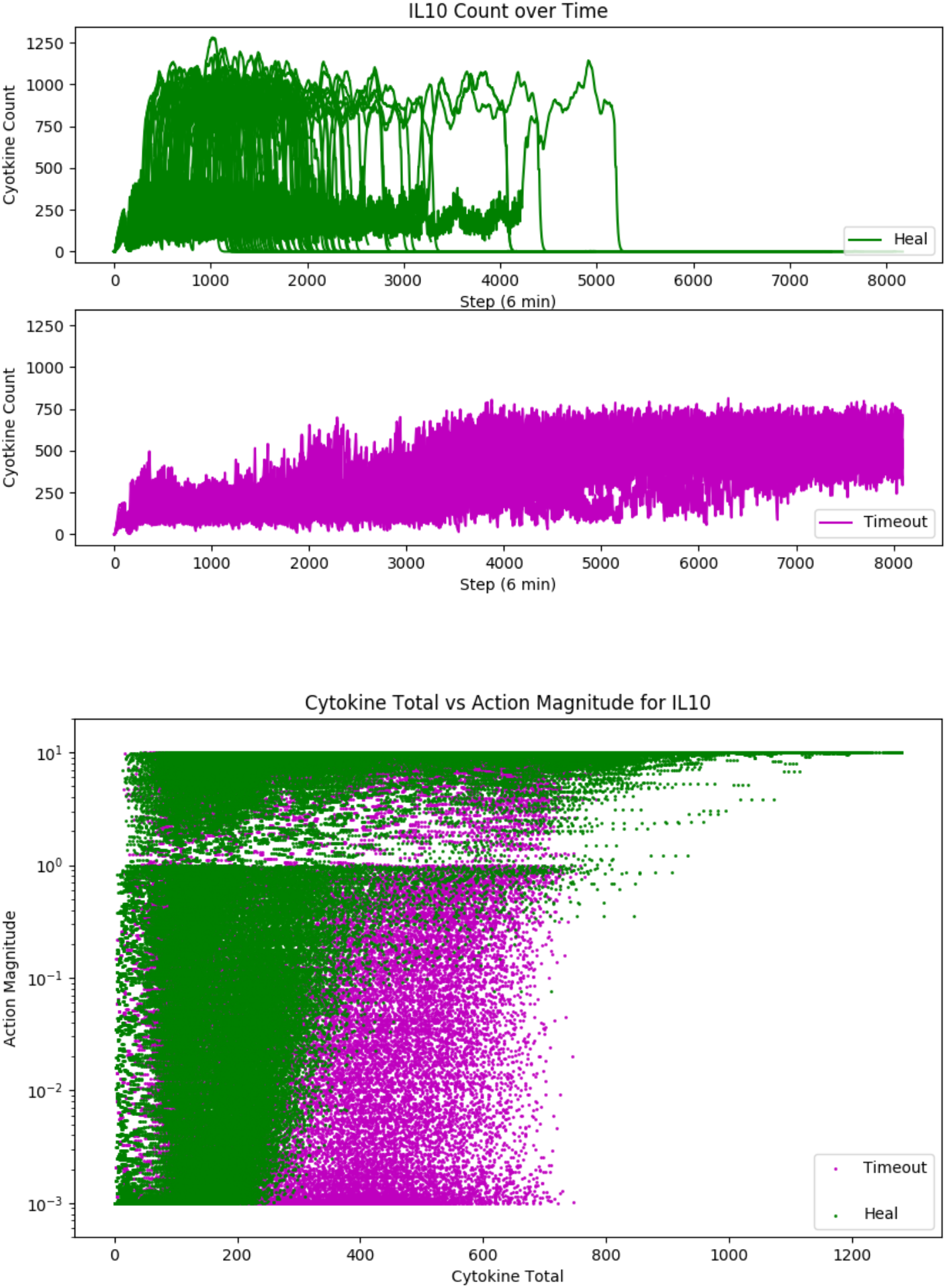
Upper panel: “Controlled” level of IL10 in Recovered simulations. Middle Panel: “Controlled” level of IL10 in Timeout/Death simulations. Lower Panel: Plot of Action Magnitude (= Control) on Y axis versus Target IL10 level on X-axis.

**Figure 9:**
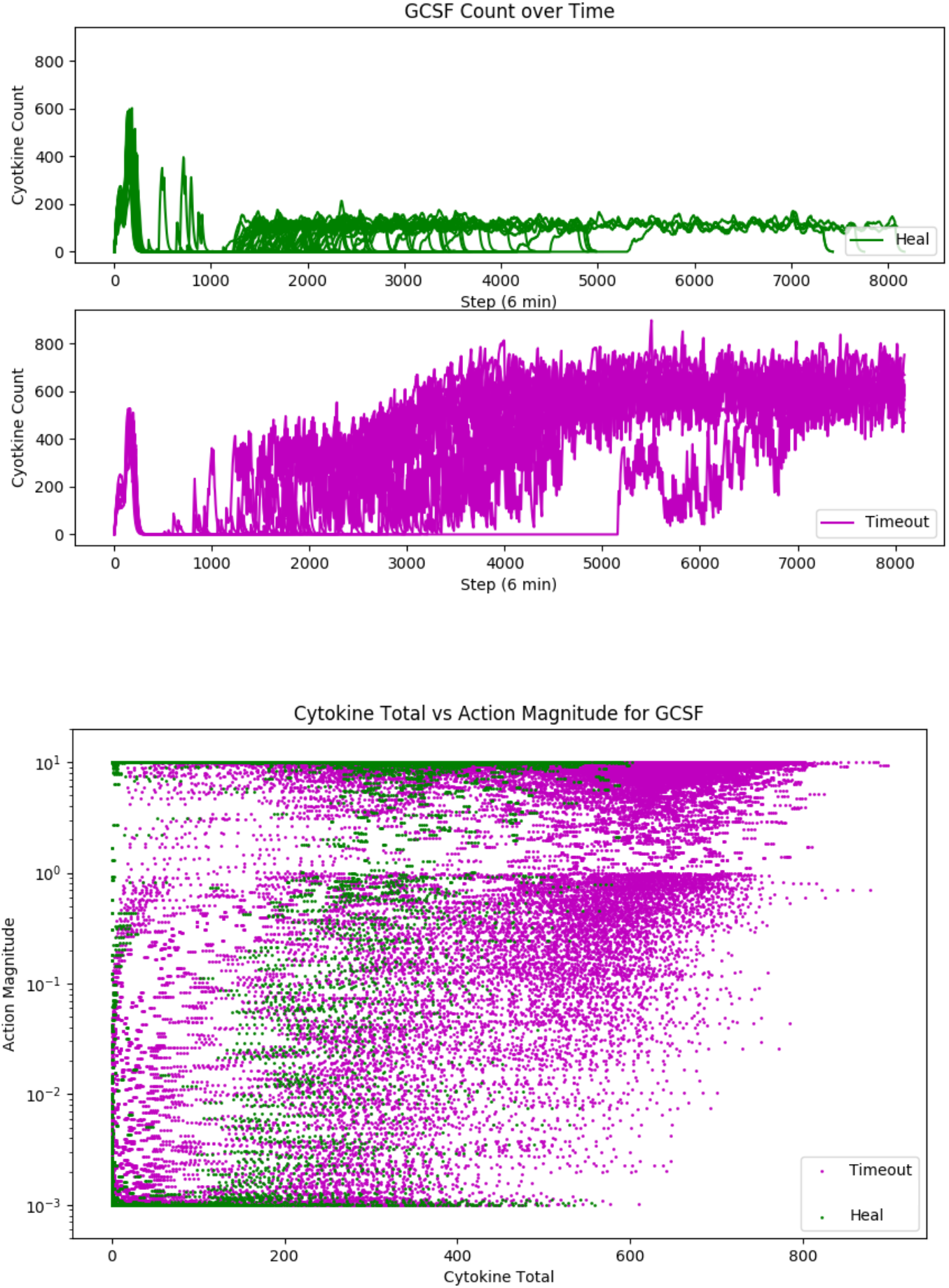
Upper panel: “Controlled” level of GCSF in Recovered simulations. Middle Panel: “Controlled” level of GCSF in Timeout/Death simulations. Lower Panel: Plot of Action Magnitude (= Control) on Y axis versus Target GCSF level on X-axis.

Of particular interest is the maintenance of above-baseline levels for IFNg seen in Figure 7; we interpret this to be due to the need for the system to maintain some degree of pro-inflammation in order to counteract the effect of microbes as per the simulation environment.

Also, with respect to the depiction of the actions taken based on mediator level present seen in the lower panels of Figure 7–9: these can be interpreted as a graphical depiction of a 1-dimensional analysis of the control policy. They are 1-dimensional insomuch that trained DRL agent is learning across the entire system state of the model, and therefore caution needs to be exercised in inferring causality in these plots. However, there is a plausible general pattern of higher levels of target mediators associated with higher magnitude of actions.

## Discussion

Sepsis has been known to involve disordered and “excessive” inflammation for half a century (25). However, attempts to modulate the inflammatory response in the face of acute infection ever since have failed to effectively translate into the clinical arena (12). COVID-19 resurrected this interest (26), with what should have been expected undecisive results. Among key lessons to be learned from the COVID-19 experience are:

1. Pathogen-agnostic disease mitigation is a critical capability in terms of readiness for future viral pandemics. While there is a certain appeal to developing viral-species specific interventions, such as anti-viral agents and vaccines, these agents have a mandatory lag-time in terms of their development; despite the impressive and unprecedented success and rapidity of COVID-19 vaccine development, it is difficult to imagine how such modalities could be made available is less than a year. Alternatively, there is a highly conserved mechanism of disease pathogenesis arising from the host inflammatory response, a shared feature of many viral infections (1–6). Developing effective strategies to control this process, while maintaining host capability to eradicate the infection, would provide a crucial capability in the early phases of any future pandemic.
2. However, the need to balance effective inflammatory/immune antimicrobial responses while mitigating the detrimental effects excessive inflammation is a highly complex task. The general failure of immunomodulation in the face of acute infection suggests that future approaches should consider this problem as complex control problem, and apply methods appropriate to solving complex control problems (13).
3. Drug repurposing is not as simple as extrapolating the putative mechanism of a drug and assuming that such a mechanism would be efficacious in a completely different context. The urgency of COVID-19 prompted the initiation of multiple potential therapies and trials based on bioplausibility; but it should be noted that every failed clinical trial presupposes that same bioplausibility. The same Translational Dilemma present in the development of new therapeutics (27) is also in play with the drug repurposing task, and requires the same readjustment of how to accomplish that task. Notably, the nature of the Translational Dilemma, i.e., the need to dynamically mechanistically-evaluate putative mechanistic bioplausibility, means that correlative approaches that utilize AI/traditional computational approaches do not provide a scientifically sound path that addresses the fundamental step in the drug evaluation process because they rely on correlative methods and the extrapolation of mechanistic-effect that has been demonstrated to be ineffective (28–30).

We have previously proposed that the integration of advanced forms of ML (specifically DRL) and high-fidelity mechanism-based simulations provides a scientifically sound path forward (13, 15). The challenges moving forward are to develop more detailed and trustworthy simulation models that can be used for training AI-controllers; these include not only being able to represent the biology in sufficient detail, but calibrating and parameterizing such models that take into account the inherent incompleteness of biological knowledge and the considerable heterogeneity seen in biological behavior (31, 32). We hope that this proof-of-concept demonstration will prompt additional investigations to improve and advance this methodology, and, critically, help drive the corresponding developments in real-time mediator/cytokine sensing and administration.

## Acknowledgements

This work was supported in part by the National Institutes of Health Award UO1EB025825 and through funding provided by the Defense Advance Research Projects Agency (DARPA) cooperative agreement HR00111950027.

